# Transforming the performance of runners with AI-driven training planning and daily adaptivity

**DOI:** 10.1101/2023.10.06.561160

**Authors:** LC Nilsson, O Södergård, J Rogestedt, CM Mattson, FJ Larsen

## Abstract

Balancing intense training with adequate recovery is key for optimal athlete performance. While insufficient or excessive training can adversely impact performance, advancements in wearable technology facilitate more effective monitoring of training and readiness level. This study combined data from Garmin watches and a 9-item questionnaire (Readiness Advisor) application to evaluate the readiness during a 39-day long training period. Two groups were studied: one with daily adaptive modifications of the training according to their readiness level and another with a static regimen where no changes were made regardless of readiness level. Results indicated that the adaptive group maintained a more consistent readiness score and showed improved physiological responses and better performance metrics. In contrast, the static group displayed non-significant improvements. This suggests that adaptive training plans, driven by individualized data and AI analytics, can significantly enhance performance outcomes and physiological adaptations for runners.

## Introduction

Athletes, elite or recreational, aiming to optimize performance face the challenge of balancing intense training stimuli with enough recovery. It is well-documented that the right exercise stimuli can enhance performance while both too little or excessive training can lead to a decline in performance. In most cases for the general population, the training load is on the low side and any increase would lead to improved performance, assuming enough rest and nutrition. The maladaptation occurring after excessive training with insufficient recovery between sessions is referred to as non-functional overreaching (NFOR), which if prolonged, can progress into the overtraining syndrome, as described by Halson and Jeukendrup (2004). In laboratory-controlled studies our research has identified that excessive training can lead to mitochondrial dysfunction, disturbances in glucose metabolism and reduced performance (Flockhart et al, 2021; Flockhart et al 2022).

With the advancements in wearable technology, monitoring training and recovery has become more straightforward, allowing continuous tracking with little disruption to an individual’s daily routine. On a broad and general level, it has been shown that the use of consumer-based wearables can lead to increase physical activity levels (Brickwood et al, 2019), and that individualized prompts generate a larger increase in physical activity than generic suggestions (Javed et al, 2023). Metrics like resting heart rate, heart rate variability, body temperature, sleep, continuous blood glucose, and subjective readiness ratings can be consistently measured and tracked. However, there is a lack of research on how to fully interpret variations in measured parameters and especially how to give recommendations and modify an athlete’s daily training based on these parameters. For instance, if an athlete experiences poorer sleep than usual, it is unclear whether reducing the training load results in better long-term outcomes than adhering to the planned program. Similarly, if all indicators suggest an athlete is well-recovered, it is uncertain whether increasing the training load leads to better outcomes. Additionally, determining the degree to which these parameters need to deviate from their baseline to warrant training adjustments remains unclear.

Based on both our published studies and ongoing research, which includes controlled laboratory-based training interventions and a continuous observational study of elite endurance athletes, we have gathered daily data on training load, performance, glucose metabolism, nervous system activation, sleep, and ratings of subjective well-being. Combined with results from physiological testing in the laboratory and real-world competition results, this comprehensive dataset has allowed us to develop and train a hybrid model combining explicit physiological relations with tools from modern AI. This model aims to differentiate between: (1) too low training loads, (2) ideal training loads, and (3) NFOR states.

In this pilot project, we utilized data from Garmin watches worn by recreational runners and combined it with feedback from the Readiness Advisor, a 9-question subjective application.

### Individuality

For each runner, we developed personalized models to analyze their training responses, a Digital Twin. This involved an ensemble method that combined time series analysis, RNNs, and regression techniques. To improve data accuracy, an LSTM network was specifically tailored to every participant’s relevant running characteristic, like heart rate, speed, and elevation profile, and used as a noise reduction module in the data parsing pipeline.. Subsequently, a daily performance metric was derived from a model assessing the speed-to-heart rate ratio. This process employed a search algorithm to find specific segments in each session. It is essential to note that the speed-to-heart rate ratio was adjusted for potential confounders. These include heart rate drift, inclination, technical terrain and recent training load.

### Optimality

Leveraging the data from the pre-processing pipeline, deep individual correlations in the response to external stimuli were modeled and a digital twin created. For each individual, we executed 5,000 simulations of various training programs to identify an optimal training load for the designated training period. Optimality was evaluated as the predicted performance by the digital twin given the hypothetically prescribed training plan.

### Adaptivity

Training schedules were recalibrated daily based on three determinants:

1. Execution of the planned training. If a runner either skipped a session or did not meet the intended load, forthcoming sessions were intensified. Conversely, if a runner exceeded the planned training load, the subsequent sessions were reduced.

2. The aggregated score from the Readiness Advisor. Should the score deviate by more than 0.5 standard deviations from an individual’s baseline in either direction, training was adjusted upwards or downwards as appropriate.

3. If the athlete reported injury or illness, the suggested training was adjusted to “Alternative Training” or “Rest Day” dependent on severity.

The primary objective of this study was to evaluate the efficacy of only the last step, adaptivity, in contrast to a non-adaptive (static) program. Thus, even the participants in the non-adaptive group received an individualized and optimal plan, based on Digital Twin simulations, for the full duration of the study period. We enrolled 20 participants, whom we randomized into two cohorts: an adaptive group receiving daily modifications to their training schedules, and a static group that, while benefiting from individualized and optimized training plans, did not receive adjustments based on training execution or daily readiness assessment but adhered to the initially generated training plan.

## Results

### Training load

The training period lasted an average of 39 days in both groups. Before the training period started the habitual number of training sessions were 3.3 +/- 0.3 sessions and 35 +/- 1 km per week in the adaptive group and 3.4 +/- 0.7 sessions and 34 +/- 2 km per week in the static group. During the training period the training increased to 4.7 +/- 0.6 sessions and 57 +/- 9 km per week in the adaptive group and 4.9 +/- 0.6 sessions and 48 +/- 8 km per week in the static group. Importantly, there were no significant differences in these parameters between groups at baseline or during the training period. See figure 1.

**Figure 1.**
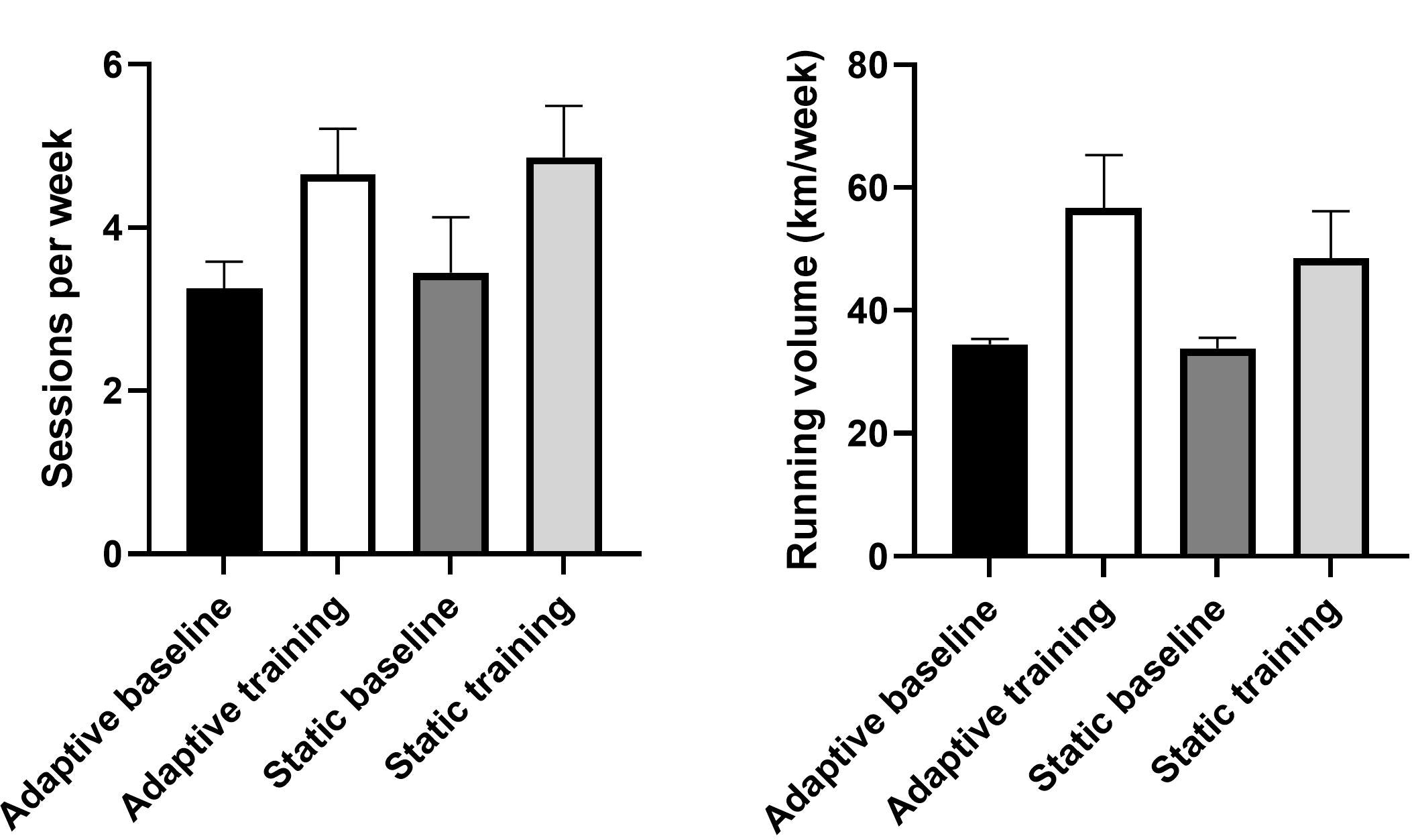
The left part of the figure indicates the average number of sessions per week and the right part illustrates the average training volume (km) per week during baseline (run-in) and during the training period in the adaptive and the static intervention groups respectively. n=7 adaptive and n=10 static.

### Adaptive training is associated with better readiness score

The weighted total score (0-100 %) of the Readiness Advisor was tracked throughout the training period. A safe zone was defined as the baseline +/- 0.5 standard deviations. The participants in the adaptive group spent in total 53 +/- 3 % of days with a score outside the safe zone. For the static group this number was significantly higher; 66 +/- 3 % of the days (t-test, p=0.01 between groups). The percentage of days with a Readiness score above the safe zone was 35 +/- 3 % in the adaptive group and 32 +/- 7 % in the static group (p=0.08, trend, non-significant), whereas the percentage of days with a lower Readiness score was 17 +/- 4 % in the adaptive group and 34 +/- 7 % in the static group p=0.08). See figure 2.

**Figure 2.**
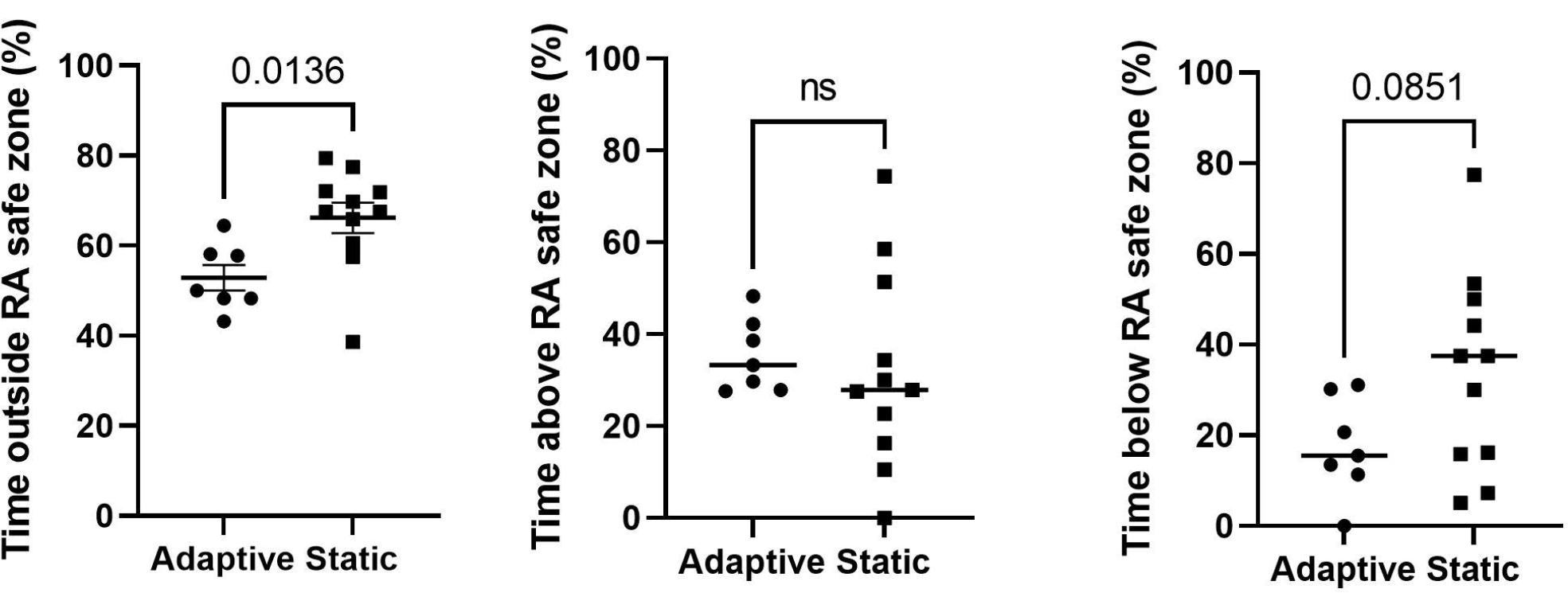
The total score (0-100%) represents a weighted sum of the 9 questions in the readiness advisor (RA). A safe zone was calculated for each individual as a baseline measure +/- 0.5 std. In the left figure, the total time (% of days) with a RA score above or below +/- 0.5 std is shown. In the middle figure the % of days with a RA score 0.5 std above baseline is shown. In the right figure the % of days with a RA score 0.5 std below baseline is shown. n=7 adaptive and n=10 static.

This means that the adaptive group spent significantly more days in the “just right” state, and that difference was mainly driven by less time with too high overall load.

### The lactate threshold is improved after adaptive training

The speed at the lactate threshold was improved from 13.3 +/- 0.4 to 13.6 +/- 0.4 km/h (p=0.02, t-test) in the adaptive group while it increased non-significantly from 13.1 +/- 0.4 to 13.3 +/- 0.5 km/h (p=0.15, t-test). Conversely, VO_2_max was unchanged from 3.55 +/- 0.17 to 3.59 +/- 0.20 l min^-1^ in the adaptive group and 3.63 +/- 0.15 to 3.57 +/- 0.18 l min^-1^ in the static group. At a submaximal running speed, ratings of perceived exertion were decreased from 12.9 +/- 0.3 to 11.3 +/- 0.5 (p=0.02, t-test) in the adaptive group while it was unchanged 11.4 +/- 0.3 to 11.6 +/- 0.3 (p=0.46, t-test) in the static group. See figure 3.

**Figure 3.**
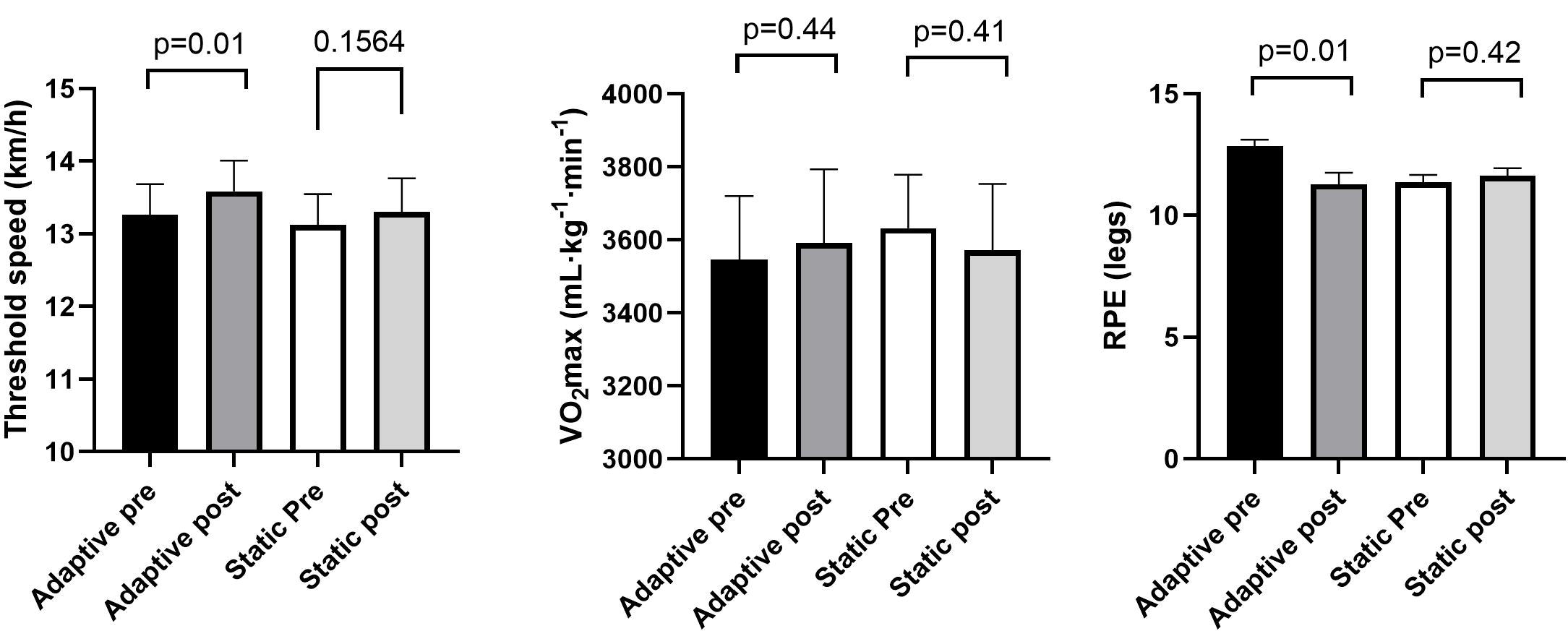
The left figure indicates the speed at the lactate threshold pre and post training in the adaptive and static group respectively. The middle figure illustrates the maximal oxygen consumption (VO_2_max) pre and post training in the adaptive and static group respectively. The right figure shows the ratings of perceived exertion (6-20) on a submaximal running speed pre and post training in the adaptive and static group respectively. n=7 adaptive and n=10 static.

### Both time to exhaustion and time-trial performance are improved after adaptive training

Time to exhaustion during the incremental max-test was improved in the adaptive group from 450 +/- 18 to 473 +/- 15 sec (p=0.03, t-test) while it was unchanged in the static group 466 +/- 18 to 466 +/- 20 sec (p=0.99, t-test). Similarly, during an outdoor 3000 meter time-trial the performance was improved in the adaptive group from 12:05 +/- 25 to 11:38 +/- 23 min:sec (p=0.01) and was improved non-significantly from 11:58 +/- 26 to 11:47 +/- 35 min:sec (p=0.21) in the static group. See figure 4.

**Figure 4.**
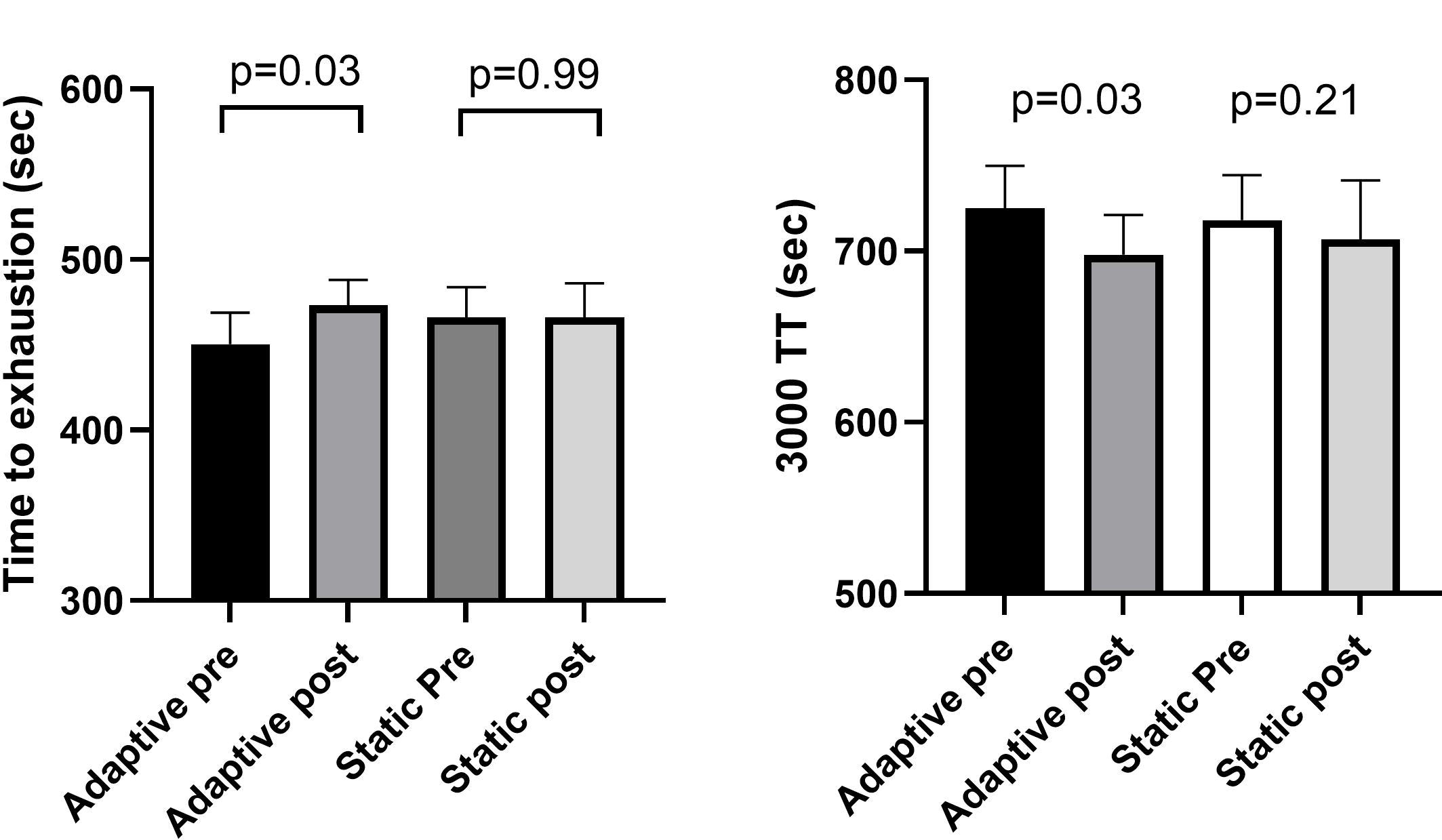
The figure to the left illustrates the time to exhaustion during the incremental running test on the treadmill pre and post training in the adaptive and static group respectively. n=7 adaptive and n=10 static. The right figure shows the finishing times in the outdoor 3000-meter time trial. n=6 adaptive and n=9 static.

Collectively, these results suggest that daily adjustment of the training is associated with a more stable readiness score, less days with low readiness, superior physiological adaptations, and better performance outcomes measured with several metrics.

## Discussion

To our knowledge, this project is the first to use a multi-dimensional approach to adjust training loads for performance optimization. While previous studies have employed heart rate variability (HRV) as a lone metric to guide training loads (see Granero-Gallegos et al, 2020 for a detailed review), they often had design limitations. For instance, the HRV and control groups did not undergo equivalent training. In numerous scenarios, an intense training protocol was assigned, and the HRV group reduced their training load if HRV metrics crossed a certain threshold. This suggests HRV-based training can mitigate overtraining, but its superiority over fixed training in enhancing performance is still up for debate. In the present project, both groups underwent equal training loads across the entire training period. A key distinction of our methodology is the model’s ability to both increase and reduce training loads based on recovery metrics, ensuring an optimal and tailored daily training regimen.

Although the overload-recovery concept is well-established and used by athletes and coaches, it is unknown what degree of overload yields the optimal performance benefits. Indeed, too small overload will not result in any positive adaptations while too great overload has been associated with negative adaptations especially on the mitochondrial level (Bellinger et al, 2020; Flockhart et al, 2021). Here we utilized a model that bidirectionally adjusted the training load when the readiness was more than 0.5 standard deviations above or below the mean.

Even in this relatively short training period of 5.5 weeks, there were distinct improvements in performance, and these improvements were superior in the adaptive intervention.

## Conclusion and Implications

The integration of advanced AI-driven methodologies to daily training adjustments based on recovery and performance metrics presents a significant stride forward in optimizing training regimens for recreational athletes. This study demonstrated the tangible benefits of an adaptive training program over a static one, with improved physiological adaptations, stable readiness scores, and superior performance outcomes for the adaptive group.

A few important observations and implications emerge from our study:

The daily adjustment of training regimens according to readiness metrics, instead of a fixed schedule, has the potential to more effectively mitigate overtraining and undertraining. This tailoring may maximize performance outcomes and overall athlete health, including maintained motivation and career longevity.

Beyond traditional metrics like heart rate variability, a multi-dimensional model that accounts for various physiological, psychological, and performance data points offers a holistic understanding of an athlete’s state. This can be crucial for devising training strategies that align with an athlete’s unique needs and responses to different stimulus.

Despite the widespread use of wearables, aiming to track both performance and health related aspects, the knowledge and use of measured data is far from what it could be. The possibilities provided by daily streams of data requires a multidimensional and interdisciplinary approach but may have the potential to generate direct and valuable suggestions to not only optimize health and performance, but also minimize the risks of illbeing, injuries and overall, negatively affected health.

The creation of digital twins for every runner illustrates the power of hyper-personalized modeling in sports science. This ushers in the scalable opportunity to move from a one-size-fits-all approach to individualized training plans based on a runner’s unique physiological response patterns.

While this study focused on recreational runners, the implications of its findings could be relevant for broader demographics. Elite athletes, fitness enthusiasts, and even individuals recovering from injuries could benefit from AI-driven adaptive programs, potentially leading to more effective training, faster recovery, and reduced injury rates.

Further research is required to validate and expand on these findings across different athletic domains, sports, and population groups. It will also be essential to analyze the long-term effects of such adaptive training methods to ascertain their implications over extended periods. In this study, deviation by more than 0.5 std was the selected level to trigger an adjustment in training load. The optimality of this selected level needs to be further studied. Collaboration with technology developers and third parties will be crucial to streamline the collection of data, and the efficiency of the AI-driven systems, ensuring they are accessible and user-friendly for athletes, trainers, and coaches.

In conclusion, the nexus of technology, AI, and sports science presents an exhilarating frontier and paradigm shift for optimizing athletic performance. As technology continues to advance and our understanding of the human body deepens, we stand on the precipice of a revolution in training methodologies that cater to the individual needs of athletes, ensuring their full potential is realized.

## Materials and Methods

### Physiological assessment

Prior to and following the training period, we evaluated baseline physiological attributes. A standardized protocol was deployed for assessing the physiological response to submaximal and maximal running. This involved a sequence of intervals, each lasting five minutes, interspersed with one-minute rest periods. During these rests, capillary blood samples were collected for lactate and glucose analysis using the Biosen C-Line Clinic (EKF-diagnostics, Barleben, Germany) and a subjective rating of perceived exertion.

The treadmill’s velocity was individually adjusted for the initial stage and subsequently increased by either 1 km/h or 2 km/h with each successive stage, continuing until a marked accumulation of blood lactate was observed. Following a brief respite, a graded maximal exertion test was initiated to determine VO_2_max. The VO_2_max value was determined as the mean value of the three highest consecutive 10-second intervals. Throughout the session, we used the Cosmed Quark CPET for real-time, breath-by-breath gas exchange monitoring. Heart rate data was recorded using participants’ personal Garmin watches. Additionally, at each stage and upon reaching exhaustion, participants were asked to indicate their perceived exertion levels utilizing the BORG scale (Borg, 1982).

On a separate day, participants were instructed to perform a 3000-meter time-trial at their best possible effort while recording the activity with their Garmin watch.

Participants: Twenty recreational runners were recruited for the study, divided into two groups: static (n=10) and adaptive (n=10). Three subjects dropped-out at random due to time-constraints or illnesses, non-related to the study protocol.

Data Collection: All participants provided historical training data from their Garmin watches, detailing their recent training patterns. This data included session duration, heart rate, running cadence, altitude change (inclination) and speed for at least 3 months prior to the study start.

The heart rate zones (1-6) were personalized based on each runner’s resting, threshold, and max heart rate as determined during the pre-tests (Mattsson CM & Larsen FJ, 2011)

Use of the Readiness Advisor (RA) App: For a period of two weeks before and throughout the training program, runners utilized the “Readiness Advisor” app. This app presented them with nine questions daily about muscle soreness, sleep, mood, energy levels, injuries, illness, food and drink intake, training load, and mental stress levels. Each parameter was rated on a scale of 0-10 using a slider. Subjects were instructed to use the app in the morning before the first training session of the day.

### Statistics

Unpaired t-tests were used to detect any changes between groups. Paired t-tests were used to detect differences between pre and post training. Normal distribution was assessed using the Shapiro-Wilk test. A p-value below 0.05 was considered to indicate statistical significance.

